# Structure basis for recognition of 26RFa by the pyroglutamylated RFamide peptide receptor

**DOI:** 10.1101/2023.12.09.570933

**Authors:** Sanshan Jin, Xin Li, Youwei Xu, Shimeng Guo, Canrong Wu, Benxun Pan, Wenwen Xin, Heng Zhang, Wen Hu, Yuling Yin, Tianwei Zhang, Kai Wu, Qingning Yuan, H. Eric Xu, Xin Xie, Yi Jiang

**Affiliations:** Lingang Laboratory, Shanghai 200031, China; State Key Laboratory of Drug Research, Shanghai Institute of Materia Medica, Chinese Academy of Sciences, Shanghai 201203, China; School of Life Science and Technology, ShanghaiTech University, Shanghai 201210, China; School of Pharmaceutical Science and Technology, Hangzhou Institute for Advanced Study, University of Chinese Academy of Sciences, Hangzhou, Zhejiang, China; University of Chinese Academy of Sciences, Beijing 100049, China; Shandong Laboratory of Yantai Drug Discovery, Bohai Rim Advanced Research Institute for Drug Discovery, Yantai, Shandong 264117, China

## Abstract

The neuropeptide 26RFa, a member of the RF-amide peptide family, activates the pyroglutamylated RF-amide peptide receptor (QRFPR), a class A GPCR. The 26RFa/QRFPR system plays critical roles in energy homeostasis, making QRFPR an attractive drug target for treating obesity, diabetes, and eating disorders. However, the lack of structural information has hindered our understanding of the peptide recognition and regulatory mechanism of QRFPR, impeding drug design efforts. In this study, we determined the cryo-EM structure of the G_q_-coupled QRFPR bound to 26RFa. The structure reveals a unique assembly mode of the receptor extracellular regions and the peptide N-terminus and elucidates the recognition mechanism of the C-terminal heptapeptide of 26RFa within the transmembrane binding pocket of QRFPR. The study also clarifies the similarities and distinctions in the binding pattern of the RF-amide moiety in five RF-amide peptides and the RY-amide segment in neuropeptide Y. These findings deepen our understanding of the RF-amide peptides recognition, aiding in the rational design of drugs targeting QRFPR and other RF-amide receptors.

---

RF-amide peptides are a family of neuropeptides widely present in most animal phyla^1,2^ which display a great sequence diversity while sharing a conserved C-terminal RF-amide sequence^1,3^. At present, five groups of RF-amide neuropeptides are identified in Mammalian. These neuropeptides include pyroglutamylated RF-amide peptide (QRFP), neuropeptide FF (NPFF), RF-amide-related peptide (RFRP or NPVF), prolactin-releasing peptide (PrRP), and kisspeptin groups. These neuropeptides act through five G protein-coupled receptors (GPCRs): pyroglutamine RF-amide peptide receptor (QRFPR, GPR103)^4^, neuropeptide FF receptor 1/2 (NPFF_1/2_R, GPR147/GPR74)^5^, prolactin-releasing peptide receptor (PrRPR, GPR10)^4^, and kisspeptin receptor (KISS1R, GPR54)^6^. These RF-amide peptides and their respective receptors play crucial roles in a wide range of neuroendocrine and behavioral functions, like modulation of feeding^7^, energy expenditure^8,9^, reproduction^10–13^, nociception^14^, and cardiovascular regulation^7^.

cDNA encoding the precursor or orthologue of 26RFa were detected in vertebrates, from fish^15^, and phasianidae^15^, to mammals^16^. In humans, 26RFa and its N-terminal extended mature peptide 43RFa are extracted from the human brain, and mRNA of QRFPR is detected in the hypothalamus, vestibular nuclei^17^, and other areas of the central nervous system. QRFPR is also detected in some peripheral organs ^18, 19^. Both 26RFa and 43RFa are endogenous ligands for QRFPR (GPR103) ^17,18^. The amino acid sequences of 26RFa are conserved among mammalian species^20^, with 26RFa from rats being slightly more potent than that from humans in calcium mobilization^21^. Upon activating by 26RFa, QPFPR couples to G_q_ and G_i/o_ proteins^18,17^ and regulates a wide range of physiological functions, such as promoting sleep in zebrafish^22^, increasing food intake^7^ and insulin sensitivity^23^, modulating aldosterone secretion^24^, regulating bone formation^19^, and nociceptive transmission^25^ in rodent. Selective agonists and antagonists of QRFPR may take effect in the treatment of metabolic imbalance (obesity, diabetes), eating disorders, and osteoporosis. The synthetic analogue of the C-terminal heptapeptide of 26RFa targeting QRFPR and have shown long-lasting orexigenic effects in mice^26, 21^, while the selective GPR103 antagonist has demonstrated obvious anorexigenic activity in vivo^27^.

Considerable efforts have been dedicated to elucidating the recognition mechanism of 26RFa by QRFPR. Previous structure-activity analyses have suggested that both the N- and C-termini of 26RFa participate in QRFPR recognition, with the C-terminus playing a particularly vital role in QRFPR activation^21^. Notably, the C-terminal heptapeptide has been identified as the minimal active segment of 26RFa^21^. This segment has served as a molecular scaffold for the development of low molecular weight peptides with enhanced potency and increased stability^21,28^. The small molecule Pyrrolo[2,3-c]pyridines, which mimic the C-terminal Arg-Phe motif of 26RFa were developed as antagonists to GPR103 ^27^. However, the absence of structural information on QRFPR has hampered our understanding of the recognition of 26RFa by QRFPR and has impeded the rational design of drugs targeting QRFPR. Furthermore, the lack of any structural information on any family members of peptide-bound RF-amide receptors has posed challenges in understanding how the conserved RF-amide group of RF-amide neuropeptides regulates their specific receptors.

In this study, we determined the structure of the 26RFa-QRFPR-G_q_ complex utilizing the cryogenic electron microscopy (cryo-EM) technique. Combined with structural and functional analysis, our findings unveil the unique assembly mode of the extracellular regions of QRFPR and the N-terminus of 26RFa. The structure also provides insights into 26RFa recognition by QRFPR and proposes a general binding pattern of RF-amide peptides.

## The overall structure of the 26RFa-QRFPR-G_q_-ScFv16 complex

To facilitate the expression of the QRFPR-G_q_ complex, we introduced a cytochrome b562RIL (BRIL) at the N-terminus of the full-length wild-type (WT) human QRFPR ^29,30^. A Gα_q_ chimera, designated as Gα_sqiN_, was engineered based on the mini-Gα_s_ scaffold with its N-terminus replaced by corresponding sequences of Gα_i1_ to facilitate the binding of scFv16. This Gα_sqiN_ chimera has been successfully used in the structure determination of GPCR-G_q_ complexes ^31–33^. Hereinafter, G_q_ refers to Gα_sqiN_ chimera unless otherwise specified.

The NanoBiT tethering strategy was employed to stabilize the QRFPR-G_q_ complex ^34^. Efficient assembly of the 26RFa-QRFPR-G_q_-scFv16 complex was achieved by incubating 26RFa with membranes from cells co-expressing the receptors, G_q_ heterotrimers, and scFv16 (Supplementary Fig. S1a). The structure of the complex was determined by cryo-EM at a global resolution of 2.73 Å (Fig. 1; Supplementary Fig. S1 and Table S1). The high-quality density map enabled precise model building for 26RFa and QRFPR containing residues E9-A346, except for two residues D249 and G258 in the intracellular loop 3 (ICL3) (Fig. 1; Supplementary Fig. S2). QRFPR displays a canonical architecture of GPCR, consisting of seven transmembrane α-helices (TM1-TM7). 26RFa almost vertically inserts into the orthosteric binding pocket of QRFPR. The C-terminal tail of 26RFa is nestled within the TM core and its N-terminus helix stretches outward and forms extensive interactions with the extracellular region of the receptor (Fig. 1).

**Fig. 1.**
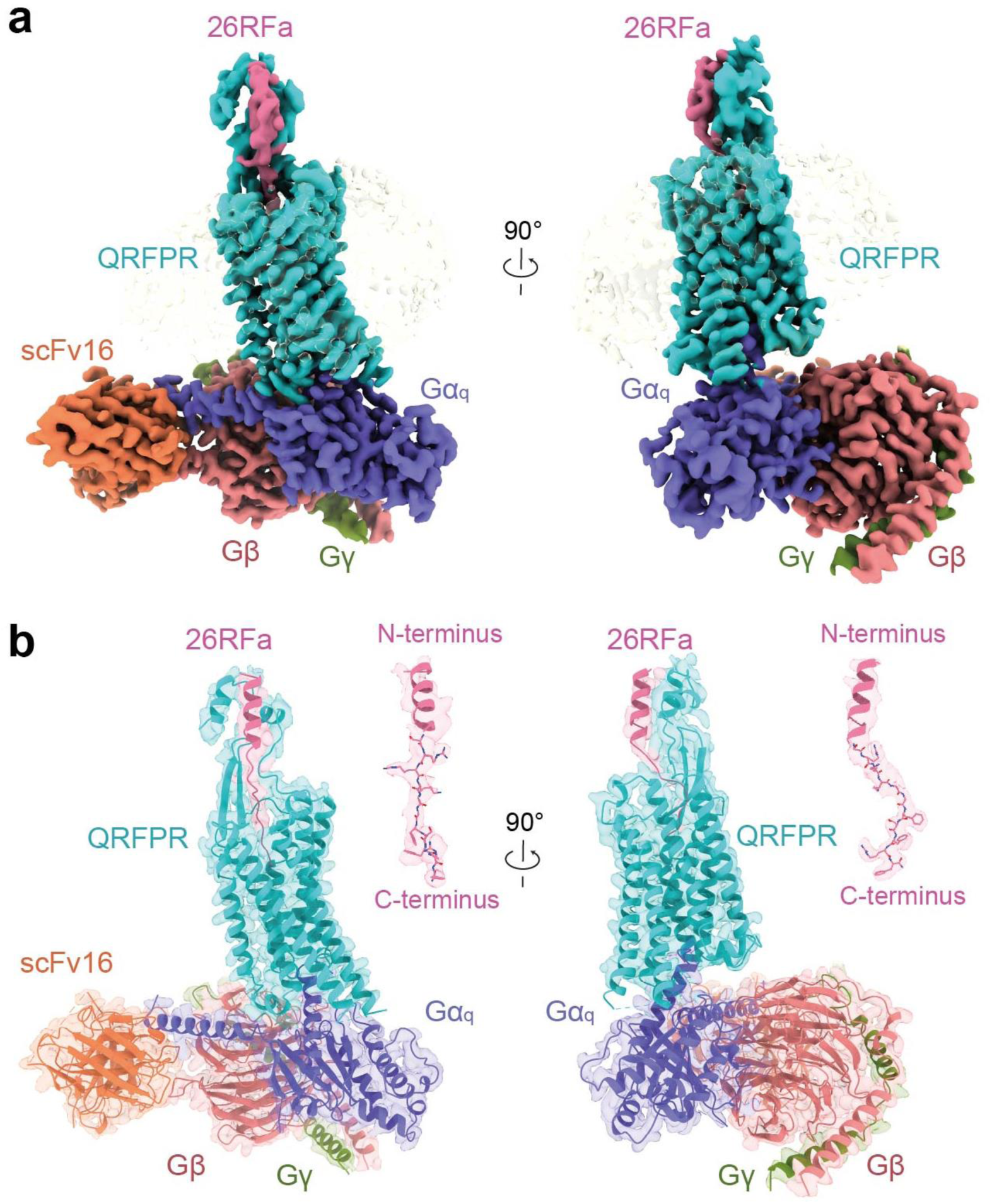
The cryo-EM structure of the 26RFa-QRFPR-G_q_-scFv16 complex. The orthogonal views of the density map (**a**) and the model (**b**) of the 26RFa-QRFPR-G_q_-scFv16 complex are shown. The components of the complex are colored as indicated.

## The unique assembly of the extracellular region of the 26RFa-QRFPR complex

Gly3-Tyr15 in 26RFa forms an α-helix, and the C terminus displays as an extended loop. Our structure of 26RFa is slightly different from that of previous NMR analysis which found the α-helical region is formed between Gly6 and Tyr15 ^35^. The N-terminus and ECL2 of QRFPR, along with the N-terminal segment of 26RFa, assemble to constitute the extracellular region of the 26RFa-QRFPR complex.

The N-terminus of QRFPR consists of two segments: a helical region (E9-R30) formed by two short perpendicular helices (E9-H18 and E23-R30) and a loop region (L31-G42) (Fig. 2a). These two short helices clip the N-terminus helix of 26RFa (Gly3-Tyr15) and cover the upper portion of the second extracellular loop (ECL2). At the top of ECL2, L193 engages in hydrophobic interactions with F11 and L34 in the receptor N-terminus and approaches the N-terminal helix of the peptide, thus bridging ECL2 to both the receptor N-terminus and the peptide (Fig. 2a). The loop region of the receptor N-terminus and the β-hairpin of ECL2 are in line with the middle loop segment of peptide (Ser16-Arg19). These elements are oriented nearly vertically to the membrane and interact with ECL1 at the transmembrane interface (Fig. 2a).

**Fig. 2.**
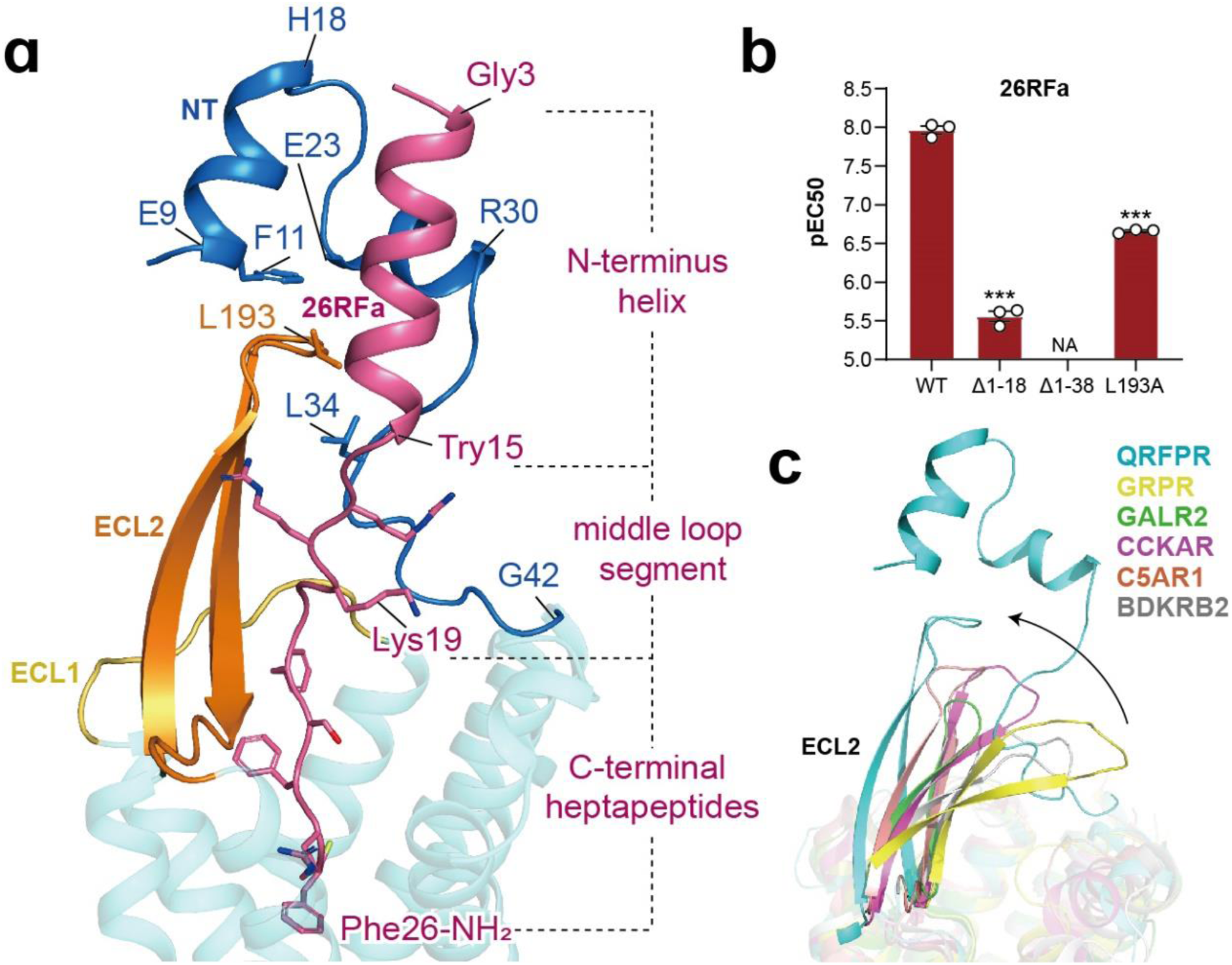
The unique assembly between the extracellular regions of QRFPR and the N-terminus of 26RFa. a, The architecture of the extracellular regions of QRFRP bound to the N-terminus of 26RFa. NT, N-terminus. **b,** Effects of mutations of the extracellular regions of QRFPR on the potency of 26RFa-induced calcium mobilization. Values are shown as mean *pEC_50_* ± S.E.M. from three independent experiments performed in triplicate. \*\*\**P*<0.001. NA, not activated. All data were analyzed by two-sided, one-way analysis of variance (ANOVA) with Tukey’s test. (D18 and D38: Δ1-18 and Δ1-38) **c,** Structure comparison of the extracellular regions of QRFPR with that of other class A peptide GPCRs. The orientation of ECL2 in QRFPR relative to other class GPCRs is depicted by a black arrow. GPRP, gastrin-releasing peptide receptor (PDB: 7W40); GALR2, galanin receptor 2 (PDB: 7WQ4); CCK_A_R, cholecystokinin A receptor (PDB: 7EZH); C5AR1, complement component 5a receptor 1 (PDB: 7Y65); BDKRB2, bradykinin receptor B2 (PDB: 7F2O).

Our functional analysis supports the crucial role of this unique extracellular assembly mode in 26RFa-induced QRFPR activity. Deleting the first short helix of the N-terminus (Δ1-18) resulted in a ∼190-fold decrease of 26RFa potency (Fig. 2b). Truncation of the entire N-terminus (Δ1-38) abolished the peptide-induced QRFPR activity, highlighting the critical role of the QRFPR N-terminus (Fig. 2b). These findings align with a previous report that removing N-terminal 12 amino acids of 26RFa, that is the helical segment clipped by the loop region of the receptor N-terminus, led to a decrease in 26RFa activity by an order of magnitude^21^. Additionally, alanine substitution of L193 at the N-terminus-ECL2-26RFa interface caused a remarkable decline in peptide activity (Fig. 2b). These structural observations and functional evidence highlight the significance of the assembly between the receptor’s extracellular region and the peptide in regulating the activity of QRFPR by 26RFa.

This assembly mode of the extracellular region in the 26RFa-QRFPR complex is unique compared to other reported class A GPCRs (Fig. 2c). In most class A GPCRs, the N-terminus is often short or lacks visible structural densities. In contrast, the N-terminus of QRFPR (E9-G42) is clearly defined in the high-quality EM density, probably due to its engagement with the N-terminus of 26RFa and ECL2 of the receptor (Fig. 2a). In addition, ECL2 in typical class A GPCRs tends to tilt towards the membrane and acts as a lid to cover the binding pocket, the ECL2 in QRFPR is vertically oriented, opening a vestibule for the binding of 26RFa (Fig. 2c).

## Recognition of the C-terminal heptapeptide by QRFPR

### Binding of Gly20-Phe24 segment within the transmembrane pocket

Besides the N-terminal helix and middle loop segment of 26RFa interacting with the extracellular regions of QRFPR, its C-terminal heptapeptide (Gly20-Phe26), the minimal active segment of 26RFa^21^, occupies the transmembrane binding pocket of QRFPR (Figs. 2a and 3a). Globally, the 26RFa segment (Gly20-Phe24) leans to TM2, but is far from TMs 5-7, leaving a large portion of unoccupied space lined by TMs 5-7 (Fig. 3a). Two glycines (Gly20, Gly21) are compactly packed into a crevice composed of residues on ECL1, ECL2, and the N-terminus (Fig. 3a). They moderately impact 26RFa activity, as substituting these two glycines of C-terminal heptapeptide with a bulkier alanine reduced the potency of 26RFa (20-26) by 4-5 folds ^21^. The side chain of Phe22 forms an intramolecular hydrophobic interaction with Phe24, which makes an extensive hydrophobic network V101^2,60^, P122^3,29^, W111^23,50^, and the conserved disulfide bond between C118^3,25^ and C201^45,50^ (Fig. 3b). Alanine substitution of W111^23,50^ and two conserved cysteines notably hamper QRFPR activation (Fig. 3d), which coincides with the non-detectable activity of the C-terminal heptapeptide when substituting Phe24 with alanine or D-Phe^21^. These results support the crucial role of Phe24 in 26RFa activity. In contrast, Ser23, whose side chain faces the spacious cavity of the peptide-binding pocket (Fig. 3a), shows a limited impact on 26RFa activity ^28^.

**Fig. 3.**
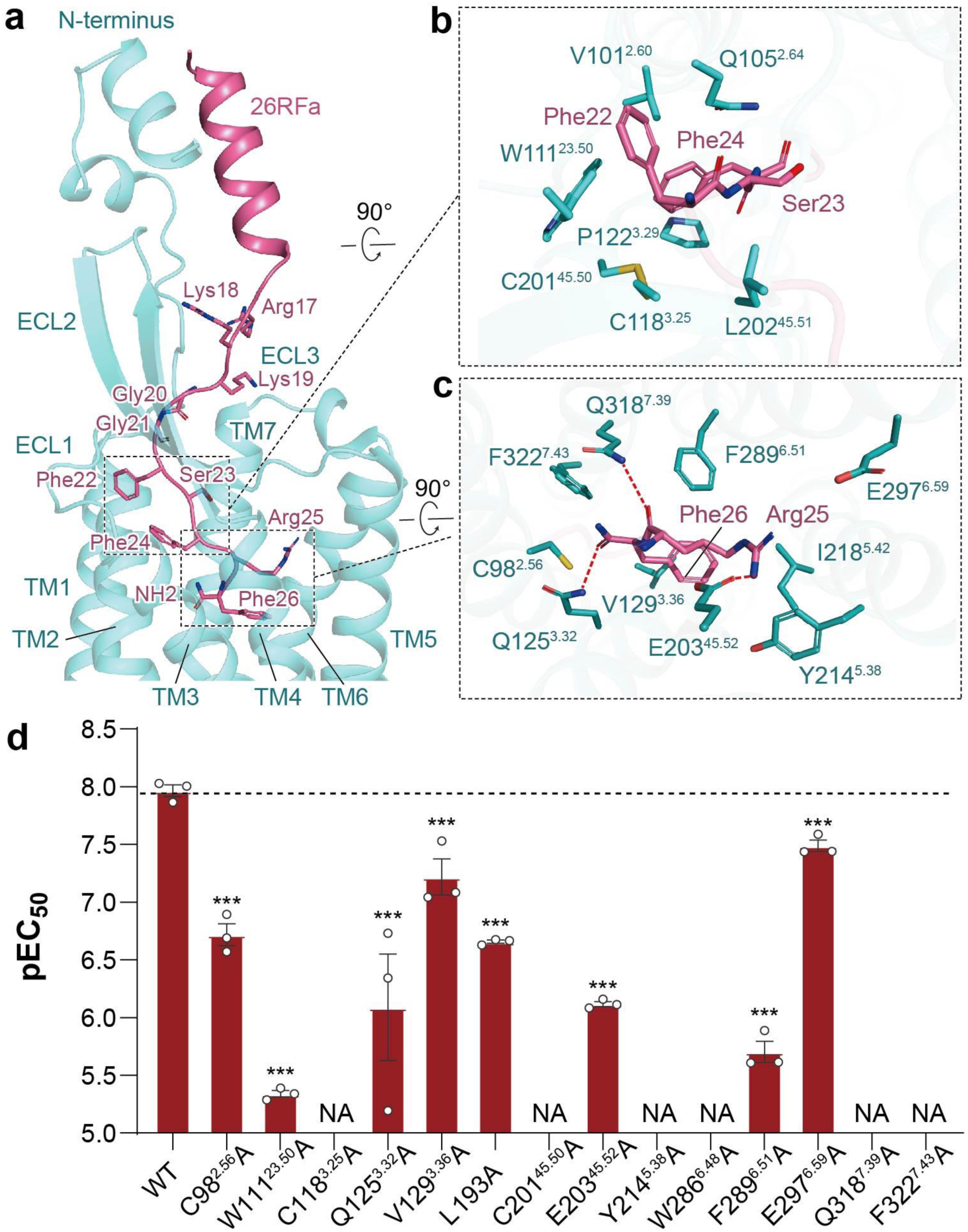
Recognition of the C-terminal heptapeptide of 26RFa by the transmembrane binding pocket of QRFPR. a, The binding pose of 26RFa on QRFPR. The heptapeptide at the C-terminus of 26RFa occupies the transmembrane binding pocket of QRFPR. **b, c,** Detail interactions between Phe22-Ser23-Phe24 (**b**) and Arg25-Phe26-amide segment (**c**) of 26RFa with residues of the transmembrane binding pocket in QRFPR. **d,** Effects of mutations in the transmembrane binding pocket of QRFRP on the potency of 26RFa-mediated calcium mobilization. Values are shown as mean *pEC_50_* ± S.E.M. from three independent experiments performed in triplicate. \*\*\**P*<0.001. NA, not activated. All data were analyzed by two-sided, one-way analysis of variance (ANOVA) with Tukey’s test.

### Recognition of the C-terminal RF-amide segment by QRFPR

The extreme C-terminus is deeply inserted into the TM helical core. The side chains of Arg25 and Phe26 point oppositely to those of Phe22 and Phe24 (Fig. 3a). The side chain of Arg25 builds a salt bridge with E203^45,52^, while its carbonyl group forms a hydrogen bond with Q318^7,39^. It also forms a cation-π interaction with Y214^5,38^ (Fig. 3c). The phenyl moiety of Phe26 is bordered by hydrophobic residues V129^3,36^, I218^5,42^, and F289^6,51^. The C-terminal amide group forms a hydrogen bond with Q125^3,22^. It is also coordinated by polar interaction with C98^2,56^ and within a distance for amide-π interaction with F322^7,43^ (Fig. 3c). All of these residues except I218^5,42^ are essential for 26RFa-induced receptor activation (Fig. 3d; Supplementary Fig. S3a), which coincides with the fact that substituting Arg25 and Phe26 with alanine abolished the 26RFa activity^21^. These findings reveal an essential role of the Arg25-Phe26-NH_2_ segment at the extreme C-terminus in 26RFa-induced QRFPR activation.

## Comparison of recognition modes of RF/RY-amide motifs by QPFR and neuropeptide Y receptors

Neuropeptide Y (NPY) and pancreatic polypeptide (PP) have a similar amidated-Arginine-Tyrosine (RY-amide) at its extreme C-terminus relative to the RF-amide motif in 26RFa (Fig. 4a). The binding pose of the N-terminus of 26RFa differs significantly from that of NPY in NPY receptors NPY_1_R, NPY_2_R (PDBs: 7X9A, 7X9B), and PP in NPY_4_R (PDB: 7X9C). The RF-amide motif in 26RFa highly overlaps with RY-amide in NPY/PP (Fig. 4b). Sequence alignment also reveals highly conserved RF-amide/RY-amide binding sites for QRFPR and NPYRs (Fig. 4c).

**Fig. 4.**
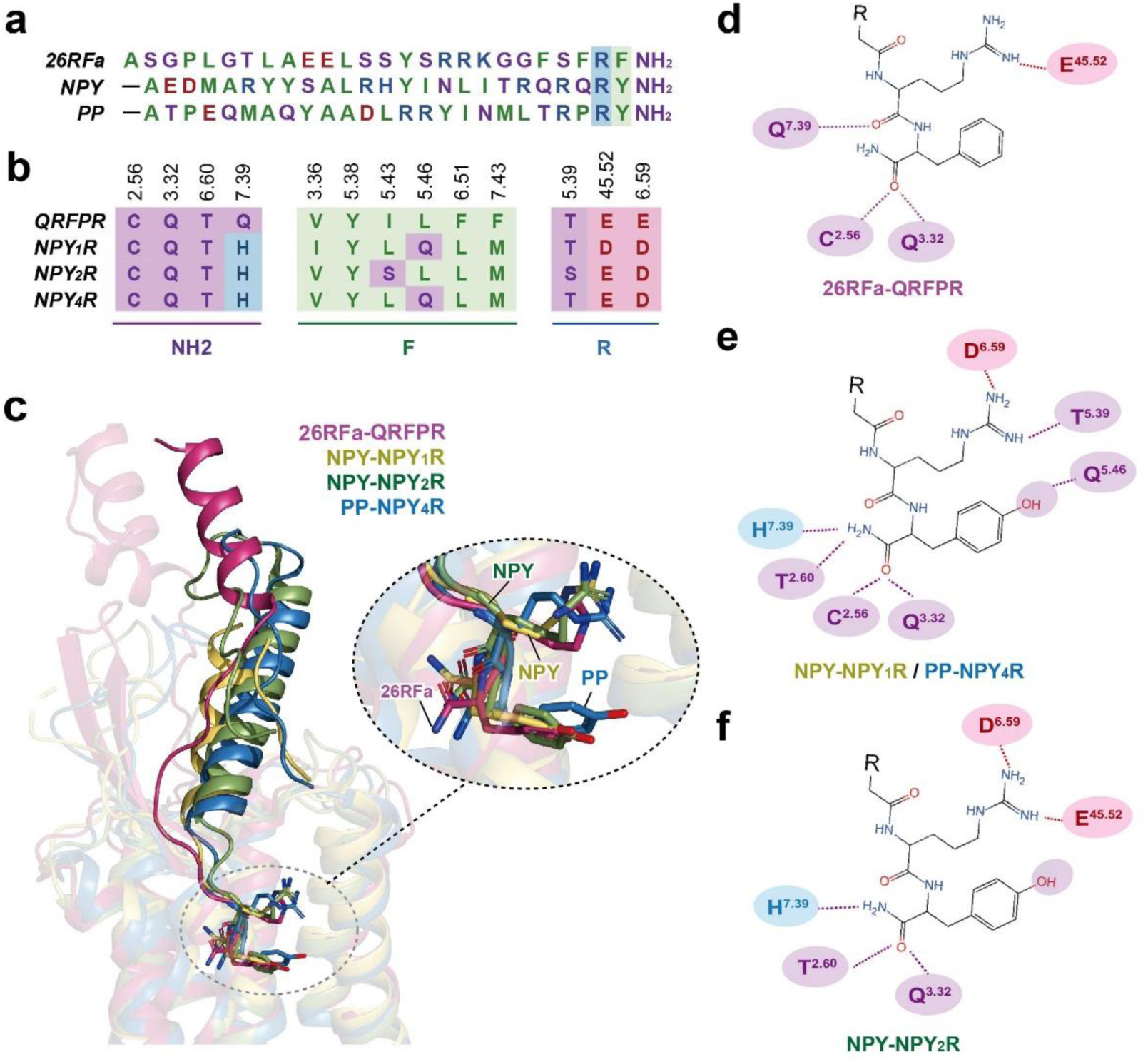
Comparison of recognition mode of peptide RF-amide and RY-amide segments by QRFPR and neuropeptide Y receptors. a, Sequence alignment of 26RFa, neuropeptide Y (NPY), and pancreatic polypeptide (PP). The N-terminal sequence of NPY and PP are omitted. These peptides show similar RF-amide and RY-amide segments at their extreme C-terminus. **b,** Sequence alignment of residues surrounding Arg25-Phe26-amide segment of 26RFa in QRFPR and cognate residues in NPY_1_R, NPY_2_R, and NPY_4_R. **c,** Structural superposition of 26RFa in QRFPR, NPY in NPY_1_R/NPY_2_R (PDB: 7X9A and 7X9B), and PP in NPY_4_R (PDB: 7X9C). **d-f**, 2D representation of key interactions between the RF-amide segment in 26RFa (**d**), RY-amide in NPY (**e**) and PP (**f**), and their specific receptors. Hydrogen bonds and salt bridges are depicted as blue- and red-dashed lines, respectively. For residues presented in **a, b, d, e,** and **f,** the light purple, blue, red, and green colors indicate polar, basic, acidic, and hydrophobic residues, respectively.

Arg25 and Arg35 in 26RFa and NPY/PP are exposed to a conserved polar environment that includes residues E^45,52^, T^5,39^, and D/E^6,59^ in QRFPR and NPYRs. However, subtle differences exist in the special binding interactions between these arginines and conserved polar residues (Fig. 4d-f). Side chains of Arg35 in NPY/PP form a conserved salt bridge with D^6,59^ and a hydrogen bond with T^5,39^ (NPY_1_R and NPY_4_R) or a salt bridge with E^45,52^ (NPY_2_R) (Fig. 4e, f). In 26RFa, Arg25 retains its salt bridge with E203^45,52^ but fails to build a salt bridge with E297^6,59^ (Fig. 3b). This is due to their greater distance resulting from an outward movement of TM6 in QRFPR relative to that of NPYRs (Supplementary Fig. S4). In addition, the conserved polar residues surrounding the amide group of 26RFa and NPY/PP, such as C^2,57^, T^2,61^, Q^3,32^, and Q/H^7,39^, participate in forming intermolecular hydrogen bonds (Fig. 4d-f). These residues are critical to 26RFa and NPY/PP activities^36^, indicating similar receptor interaction modes of the amide group between 26RFa and NPY/PP.

Phe26 in 26RFa shows distinct binding characteristics compared with Tyr36 for QRFPR and NPYRs. The hydroxyl group of Tyr36 in NPY_1_R and NPY_4_R forms hydrogen bonds with Q^5,46^, an interaction crucial for NPY/PP activity^36^ (Fig. 4e). However, in NPY_2_R, this hydrogen bond is absent due to the substitution of glutamine with leucine at position 5.46, which lacks a hydrogen bond donor (Fig. 4c, f). Similarly, QRFPR also has a leucine at position 5.46, which cannot form a corresponding hydrogen bond with Phe26 in 26RFa. Furthermore, in contrast to the notable impact of Q^5,46^ on the NPY/PP activity for NPY_1_R and NPY_4_R, the presence of L^5,46^ in QRFPR and NPY_2_R shows negligible effects on the activities of the specific peptide^36^ (Supplementary Fig. S3). These findings suggest that Phe26 and Tyr36 in 26RFa and NPY/PP have distinct roles in activating their specific target receptors.

## Comparison of RF-amide recognition mode among RF-amide peptides

The RF-amide C-terminal extremity is a molecular signature of 26RFa and other RF-amide peptides, such as NPFF, NPVF, PrRP, and kisspeptin (Fig. 5a). It is believed that the RF-amide segments within these peptides play a pivotal role in determining their bioactivity ^37^. The RF-amide-interacting residues in QRFPR are relatively conserved across RF-amide peptide receptors (Fig. 5b). This raises the intriguing question of whether these RF-amide segments exhibit a common interaction pattern with their specific receptors. We thus performed mutagenesis and functional analyses to investigate the contribution of critical residues in 26RFa to the activities of other RF-amide peptides.

**Fig. 5.**
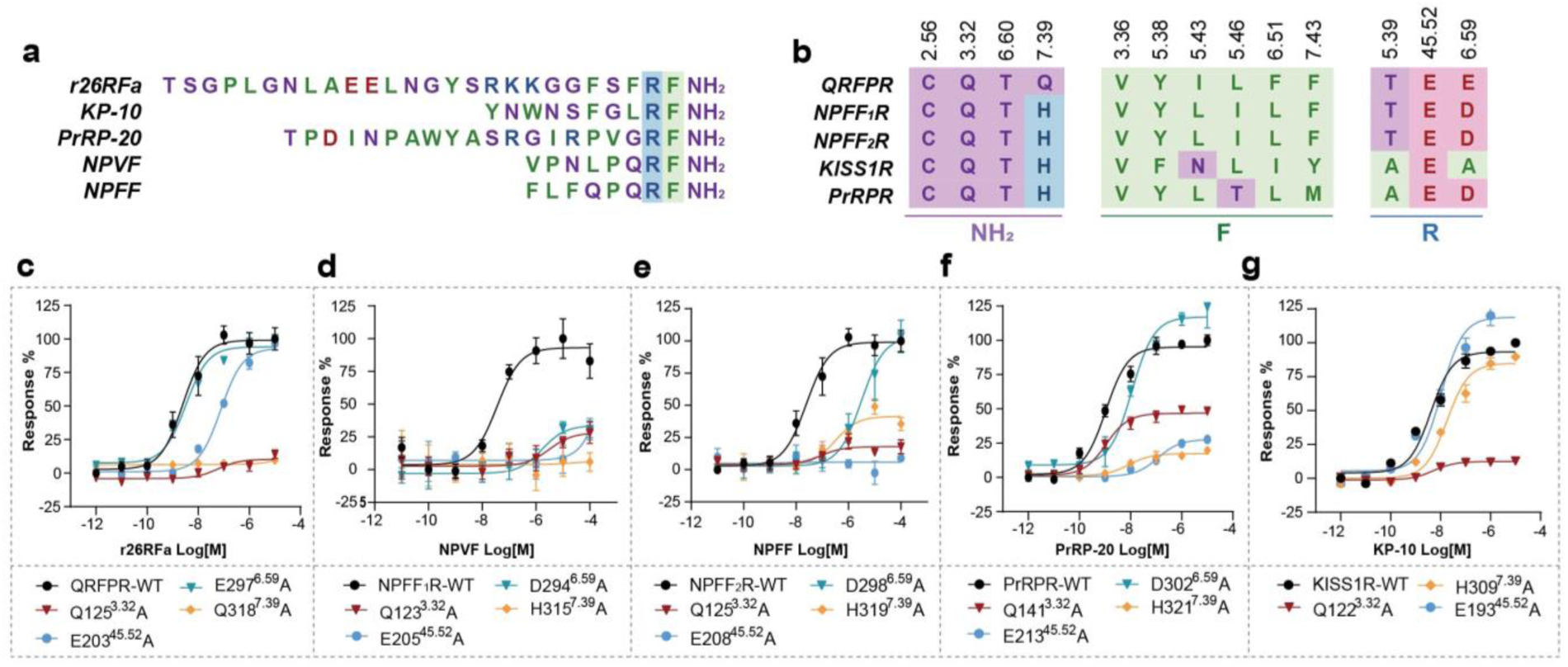
Comparison of RF-amide segment of RF-amide peptides recognized by their specific receptors. a, Sequence alignment of RF-amide peptides, which shares a conserved C-terminal RF-amide segment. **b,** Sequence alignment of RF-amide-interacting residues across RF-amide peptide receptors. For residues presented in **a** and **b,** the light purple, blue, red, and green colors indicate polar, basic, acidic, and hydrophobic residues, respectively. **c-g**, Effects of mutation in the RF-amide binding pocket on calcium mobilization. **a,** 26RFa/QRFPR; **b,** NPVF/NPFF_1_R; **c,** NPFF/NPFF_2_R; **d,** PrRP/PrRPR; **e,** KP-10/KISS1R.

In a previous study, D^6,59^ has been found important for the activation potency of NPFF1/2R^38^ and PrPR^39^. Supported by our structure and functional data, the arginine in the RF-amide segment may form salt bridges with highly conserved D/E^6,59^ and E^45,52^ across RF-amide peptide receptors (Fig. 3c). The residue D/E^6,59^ is critical for the activities of NPVF, NPFF. It also shows a moderate impact on PrRP activity but almost does not affect 26RFa activity (Fig. 5c-f). Interestingly, in KISS1R, the positively charged D/E^6,59^ is replaced with a alanine (Fig. 5b), which is unable to form a salt bridge with the arginine in the RF-amide segment of KP-10, the smallest highly potent segment of kisspeptin^40,41^. As for E^45,52^, its substitution with alanine resulted in a remarkable decline of peptide activity or potency of 26RFa, NPVF, NPFF, and PrRP towards QRFPR, NPFF_1_R, NPFF_2_R, and PrRPR, respectively (Fig. 5c-f). However, this mutation has a much weaker impact on KP-10 activity for KISS1R (Fig. 5g).

The phenylalanine in the RF-amide segment of 26RFa forms hydrophobic interactions with V129^3,36^ and F289^6,51^ in QRFPR, both of which substantially contribute to 26RFa activity (Fig. 3c, d). The valine at position 3.36 is identical, whereas the F289^6,51^ in QPFPR is substituted with leucine in other RF-amide receptors (Fig. 5b). The alanine mutation of F289^6,51^ resulted in a dramatic decrease in the activity of 26RFa (Fig. 3d), while a leucine substitution does not impact the peptide activity (Supplementary Fig. S3a). These findings highlight the significance of the hydrophobic interaction between Phe26 and F289^6,51^ for the activity of 26RFa and also suggest a potential role for V^3,36^ and L^6,51^ in the peptide activity towards other RF-amide receptors. Q123^3,32^ and Q318^7,39^ surrounding the amide group form intermolecular hydrogen bonds with the RF-amide segment in 26RFa, which are essential for the peptide’s activity (Fig. 3c, d). Sequence alignment shows that the glutamine at position 3.32 is conserved, whereas glutamine at position 7.39 in QRFPR is substituted with histidine in other RF-amide receptors (Fig. 5b). Both Q^3,32^ and Q/H^7,39^ play critical roles in the activity of 26RFa, NPVF, NPFF, and PrRP (Fig. 5c-f). In the case of KP-10, Q^3,32^ also exhibits a remarkable impact. Differently, H^7,39^ shows only minor effects on its activity (Fig. 5g).

Collectively, in contrast to KP-10, the RF-amide segment is more crucial for the activity of other RF-amide peptides. This finding is supported by the fact that, in contrast to KP-10, the alanine substitution of arginine had a more significant impact on the activity of other RF-amide peptides^37,42^.

## Activation mechanism of QRFPR

Structural comparison of the 26RFa-QRFPR complex with its homologous receptor NPY1R in inactive and active states (PDB: 5ZBQ and PDB: 7X9A) reveals a notable activation feature of QRFPR. Akin to the active NPY1R, the cytoplasmic end of TM6 of QRFPR undergoes a pronounced outward displacement in contrast to the inactive NPY1R, the hallmark of class A GPCR activation. Concomitantly, TM7 of QRFPR shows a slightly inward shift toward the core of the helical bundle (Supplementary Fig. S5a, b). These structural observations support the fact that the 26RFa bound QRFPR is indeed in the active state.

Similar to the tyrosine in the RY-amide segment of NPY, the phenyl group of phenylalanine in the RF-amide moiety of 26RFa inserts into the helical core, enabling direct contact with W286^6,48^ (Supplementary Fig. S5c), the conserved toggle switch residue in class A GPCRs responsible for peptide-induced receptor activation^43,44^. The insertion may lead to a rotameric switch of W^6,48^ and induce the rearrangement of the residues in other “micro-switch” motifs, including P^5,50^T^3,40^F^6,44^, N^7,49^P^7,50^xxY^7,53^, and E^3,49^R^3,50^Y^3,51^ (Supplementary Fig. S5d-f). These conformational changes are largely similar to that in the active NPY_1_R, indicating a shared activation mechanism between QRFPR and NPY_1_R.

## Conclusions

Our structure reveals that the 26RFa-QRFPR complex has a unique assembly mode of the extracellular regions of QRFPR and the N-terminal of 26RFa, which differs from other class A GPCRs with reported structures. The unique organization of the complex is crucial for the activation of QRFPR by 26RFa, as supported by our truncation analysis on QRFPR and the observed gradual decline in bioactivity for N-terminal consecutive truncated 26RFa analogues.

The C-terminal heptapeptide in 26RFa, which is evolutionarily conserved, displays a comparable activity to that of full-length 26RFa. This heptapeptide establishes an interaction network with residues in the transmembrane (TM) binding pocket. The low occupancy of the Gly20-Phe24 segment in the transmembrane binding pocket results in Ser23 being exposed to a spacious area without significant interactions with neighboring residues in QRFPR. This binding mode presents an opportunity for optimizing peptide activity. For instance, substituting Ser23 with the bulkier norvaline led to a peptide analogue that was 3-fold more potent than the native C-terminal heptapeptide of 26RFa^21^. Furthermore, our structures elucidate the recognition mechanism of the RF-amide segment at the extreme C-terminus of 26RFa by QRFPR within the transmembrane binding pocket.

The structural comparison reveals that the RF-amide moiety of 26RFa shares a similar binding mode with the RY-amide group of NPY for their specific receptors. Our sequence alignment and mutagenesis analyses also suggest largely conserved recognition modes of RF-amide peptides. Therefore, we propose a general binding pattern for RF-amide peptides in QRFPR, NPFF_1/2_R, and PrRPR, albeit with some distinctions in the case of KISS1R. In summary, our structural findings provide insights into the mechanisms of peptide recognition and activation of QRFPR and propose a similar binding pattern for the RF-amide segment of RF-amide peptides across their specific receptors. These discoveries offer opportunities for the design of QRFPR-targeting drugs.

## Methods

### Constructs

The human wild-type (WT) QRFPR gene was codon-optimized for insect cell (Hi5) expression. Full-length optimized QRFPR sequence was cloned into a modified pFastBac vector (Thermo Fisher) with a thermally stabilized N-terminal BRIL^29^ tag facilitating receptor expression and a C-terminal 8×His tag for receptor affinity purification. NanoBiT was split into a large subunit (LgBiT, 18 kDa) and a complementary small subunit (SmBiT, peptide, VTGYRLFEEIL). A homolog peptide of SmBiT, HiBiT (peptide 86, VSGWRLFKKIS), shows a high affinity for LgBiT^45^. We fused the LgBiT to the C-terminal of the receptor and the HiBiT with a 15-amino acid (15AA) polypeptide linker (GSSGGGGSGGGGSSG) at the C-terminus of Rat Gβ1. This NanoBiT tethering strategy^34^ was introduced to improve the structural homogeneity and stability of the QRFPR-G protein complex. The engineered Gα_q_ was designed based on the mini-Gα_s_ skeleton ^46^. 18 amino acids from the N-termini of Gα_q_ were replaced by the counterpart of the Gα_i_ which was responsible for scFv16 binding^47^. Two dominant-negative mutations (corresponding to G203A and A326S)^48^ were incorporated to decrease the affinity of nucleotide-binding to stabilize the Gαβγ complex. The engineered Gα_q_, Gβ1, bovine Gγ2, and scFv16 were cloned into the pFastBac vector respectively.

### Expression and purification of complexes

Bac-to-Bac Baculovirus was introduced in our expression system. Baculoviruses of FLAG-Bril-QRFPR (1-431)-LgBiT-H8, engineered Gα_q_, Gβ1-15AA-HiBiT, Gγ2, and scFv16 co-infected Hight Five (Hi5) insect cells (Invitrogen) in logarithmic phase at a 1:1:1:1:1 ratio. Hi5 cells were cultured in SIM-HF (SinoBiological) serum-free medium. Scaled-up insect cell grown in suspension at 27 °C and rotating at 120 rpm then harvested after 48 h and stored at -80 for use.

The frozen cells were thawed at 37°C and resuspended in lysis buffer containing 20 mM HEPES, pH 7.5, 150 mM NaCl, 10% (v/v) glycerol, and EDTA-free protease inhibitor cocktail (TargetMol) and lysed by dounce. In the homogenization stage, 25 mU/mL Apyrase (Sigma-Aldrich) and 10 μM rat 26RFa (GenScript) were added into the cell lysis and incubated at room temperature (RT) for 1.5 h. The cell lysis was then solubilized in 0.5% (w/v) lauryl maltose neopentylglycol (LMNG, Anatrace), and 0.1% (w/v) cholesterol hemisuccinate (CHS, Anatrace) for 2 h at 4°C. The supernatant was collected after centrifugation at 65,000 g for 35 min and was incubated with TALON^®^ Metal Affinity Resin (TaKaRa) for 2 h at 4°C. The resin was packed and washed with 30 column volumes of wash buffer (20 mM HEPES pH 7.5, 150 mM NaCl, 0.01% (w/v) LMNG, and 0.002% CHS, 10 mM imidazole, 1 μM rat 26RFa) and finally eluted in the buffer containing 300 mM imidazole. The eluted sample was concentrated using an Amicon Ultra Centrifugal Filter (MWCO 100 kDa) and purified by Superose 6 Increase 10/300 column (GE Healthcare) pre-equilibrated with buffer containing 20 mM HEPES pH 7.5, 150 mM NaCl, 0.00075% (w/v) LMNG, 0.00025% (w/v) GDN (Anatrace) and 0.00015% CHS, 1 μM 26RFa were used). Peak fractions were collected and concentrated for the cryo-EM study.

### Cryo-EM data collection

Cryo-EM grids were prepared with the Vitrobot Mark IV plunger (FEI) set to 6°C and 100% humidity. Three microliters of the sample were applied to the glow-discharged gold R1.2/1.3 holey carbon grids. The sample was incubated for 10 s on the grids before blotting for 4.5 s (double-sided, blot force 1) and flash-frozen in liquid ethane immediately. For 26RF-QRFPR-Gq complex datasets, 3,657 movies were collected respectively on a Titan Krios equipped with a Falcon 4 direct electron detection device at 300 kV with a magnification of 165,000, corresponding to a pixel size 0.73 Å. Image acquisition was performed with EPU Software (FEI Eindhoven, Netherlands). We collected a total of 36 frames accumulating to a total dose of 50 e-Å^-2^ over 2.5 s exposure.

### Cryo-EM image processing

MotionCor2 was used to perform the frame-based motion-correction algorithm to generate a drift-corrected micrograph for further processing ^49,50^. All subsequent steps including contrast transfer function (CTF) estimation, particle picking and extraction, two-dimensional (2D) classification, ab-initio reconstruction, hetero refinement, non-uniform refinement, local refinement, and local resolution estimation were performed using cryoSPARC^51^.

For the dataset, 3,657 dose-weighted micrographs were imported into cryoSPARC, and CTF parameters were estimated using patch-CTF. A blob picker was used for initial particle selection from a few micrographs followed by 2D classification to generate good templates. Subsequently, 2,397,272 particles were picked by template picker from the full set of micrographs and extracted using a pixel size of 2.92 Å. After two rounds of 2D classification, some of the classes that showed bad features were selected to generate three bad references by ab-initio reconstruction. And emd_32313^52^ map was imported as a good reference. Using these references, the full set of particles was subjected to four rounds of heterogeneous refinement, resulting in a 3.32 Å map reconstructed from 99,325 particles. 20 classes of 2D templates were generated from the 3D maps. Subsequently, 2,449,719 particles were picked by template picker from the full set of micrographs and extracted using a pixel size of 2.92 Å. After one round of 2D classification, 1,850,060 particles were selected for further heterogenous refinement. After three rounds of heterogeneous refinement, 393,048 particles remained, and re-extraction using a pixel size of 0.73Å. Following non-uniform refinement and local refinement, a map reached a resolution of 2.88 Å. Several rounds of heterogeneous refinement were conducted with updated reference maps, and 202,884 particles remained. Following non-uniform refinement and local refinement, the final map reached a resolution of 2.73 Å. We also performed postprocessing of the final maps with DeepEMhancer^53^

### Model building

A predicted QRFPR structure from Alphafold2 was used as the starting reference model for receptor building^54^. Structures of Gα_q_, Gβ, Gγ, and the scFv16 were derived from PDB entry 8IUL ^55^ and were rigid body fit into the density. All models were fitted into the EM density map using UCSF Chimera^56^ followed by iterative rounds of manual adjustment and automated rebuilding in COOT^57^ and PHENIX^58^, respectively. The model was finalized by rebuilding in ISOLDE^59^ followed by refinement in PHENIX with torsion-angle restraints to the input model. The final model statistics were validated using Comprehensive validation (cryo-EM) in PHENIX^58^ and provided in the Supplementary Table S1. All structural figures were prepared using Chimera ^56^, Chimera X^60^, and PyMOL (Schrödinger, LLC.).

## Functional assay

### Plasmids construct for functional assay

The wild-type QRFPR, PrPRR, KISS1R, NPFF_1_R, or NPFF_2_R gene was subcloned into the pcDNA3.0 vector with the addition of an N-terminal HA tag. All of the mutations used for functional studies were generated by QuickChange PCR and were verified by DNA sequencing.

## Cell transfection

HEK293 cells were obtained from ATCC (Manassas, VA, USA) and cultured in DMEM supplemented with 10% (v/v) FBS, 100 mg/L penicillin, and 100 mg/L streptomycin in 5% CO_2_ at 37℃. For transient transfection, approximately 2.5×10^6^ cells were mixed with 2 µg plasmids in 200 µL transfection buffer, and electroporation was carried out with a Scientz-2C electroporation apparatus (Scientz Biotech, Ningbo, China). The experiments were carried out 24 hours after transfection.

## Calcium Mobilization Assay

For QRFPR, PrRPR, and KISS1R, HEK293 transfected with WT receptor or its mutations were seeded at a density of 4×10^4^ per well into 96-well culture plates and incubated for 24 hours at 37℃ in 5% CO_2_. For NPFF_1_R and NPFF_2_R, plasmids were transfected into HEK293 stably expressing Gα_16_. The next day, Cells were incubated with 2 μmol/L Fluo-4 AM in HBSS supplemented with 5.6 mmol/L D-glucose and 250 μmol/L sulfinpyrazone at 37°C for 45 min. After washing, cells were added with 50 μL HBSS and incubated at room temperature for 10 min, then 25 μL agonist buffer was dispensed into the well using a FlexStation III microplate reader (Molecular Devices), and intracellular calcium change was recorded at an excitation wavelength of 485 nm and an emission wavelength of 525 nm. EC_50_ and Emax values for each curve were calculated by Prism 8.0 software (GraphPad Software).

## Surface expression analysis

24 hours after transfection, cells were washed with PBS, fixed with 4% PFA for 15 min, and then blocked with 2% BSA for 1 h. Next, cells were incubated with the polyclonal anti-HA (Sigma, H6908) overnight at 4°C and then horseradish peroxidase (HRP)-conjugated anti-rabbit antibody (Cell Signaling Technology, 7074S) for 1 h at room temperature. Then cells were washed and incubated with 50 μL tetramethylbenzidine (Sigma, T0440) for 30 min before the reaction was stopped with 25 μL TMB Substrate stop solution (Beyotime, P0215). Absorbance at 450 nm was quantified using a FlexStation III microplate reader (Molecular Devices).

## Acknowledgments

The cryo-EM data were collected at the Advanced Center for Electron Microscopy, Shanghai Institute of Materia Medica (SIMM). We thank all staff at the institution for their assistance in cryo-EM data collection. This work was partially supported by the Lingang Laboratory, Grant No. LG-GG-202204-01 (Y.J. and H.E.X.); the National Natural Science Foundation (32171187 to Y.J., 82121005 to X.X, Y.J. and H.E.X., 82330113 to X.X., 32130022 to H.E.X., and 82304579 to S.G.), CAS Strategic Priority Research Program (XDB37030103 to H.E.X.); Shanghai Municipal Science and Technology Major Project (2019SHZDZX02 to H.E.X.); Shanghai Municipal Science and Technology Major Project (H.E.X.); the National Key R&D Program of China (2018YFA0507002 to H.E.X.).

## Author contributions

S.S.J. screened and optimized the expression constructs and purification conditions, prepared the protein samples for final structure determination, participated in cryo-EM grid inspection and data collection, analyzed the structures, and prepared the figures and wrote the initial manuscript; X.L. and S.M.G. prepared the constructs for functional assays, conducted functional studies and functional data analysis, W.W.X. performed some of the functional assays. C.R.W. and Y.W.X. performed cryo-EM grid preparation, cryo-EM grid inspection and data collection, structure determination, built and refined the structure model, and participated in protein sample optimization, figure and manuscript preparation; B.X.P., H.Z., W.H., Y.Y.L., T.W.Z., K.W. and Q.N.Y. participated in data collection and analysis; H.E.X. conceived and supervised the project; X.X. conceived and supervised the functional study. Y.J. initiated, conceived and supervised the project, analyzed the structure, prepared the figures and wrote the manuscript with input from all co-authors.

## Data availability

All data is available in the main text or the supplementary materials. Materials are available from the corresponding authors upon reasonable request.

---

**Supplementary Fig. S1.**
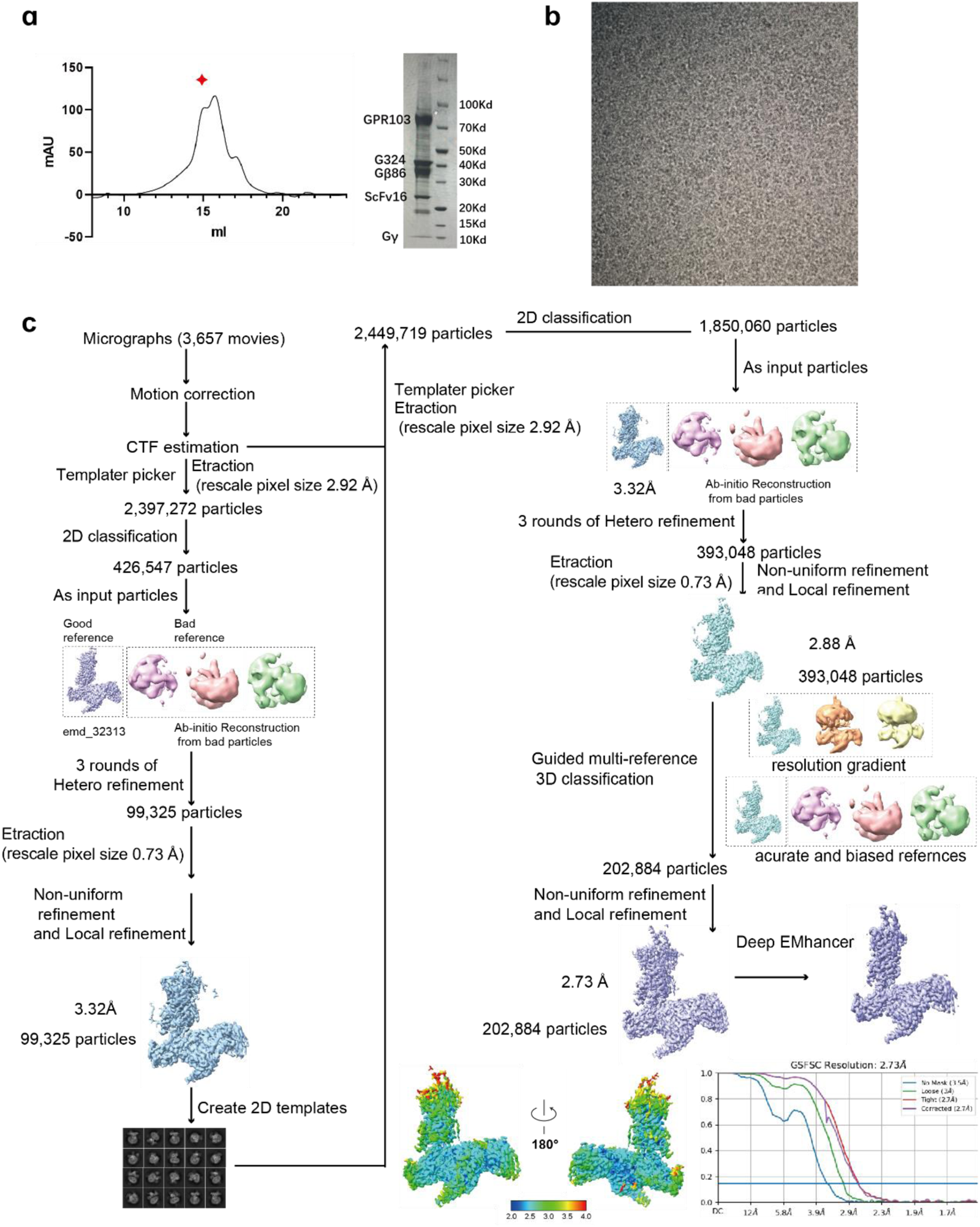
Purification and cryo-EM data processing of the 26RF-QRFPR-G_q-_ scFV16 complex. **a**, Representative size-exclusion chromatography elution profile and SDS-PAGE analysis of the 26RF-QRFPR-G_q_-scFV16 complex. The elution peak of the complex monomer is indicated by a red star. **b**, Representative micrographs of the 26RF-QRFPR-G_q_-scFV16 complex from CTF estimation. **c**, The flowchart of data processing for the 26RF-QRFPR-G_q_-scFV16 complex. Detailed descriptions can be found in the Methods section.

**Supplementary Fig. S2.**
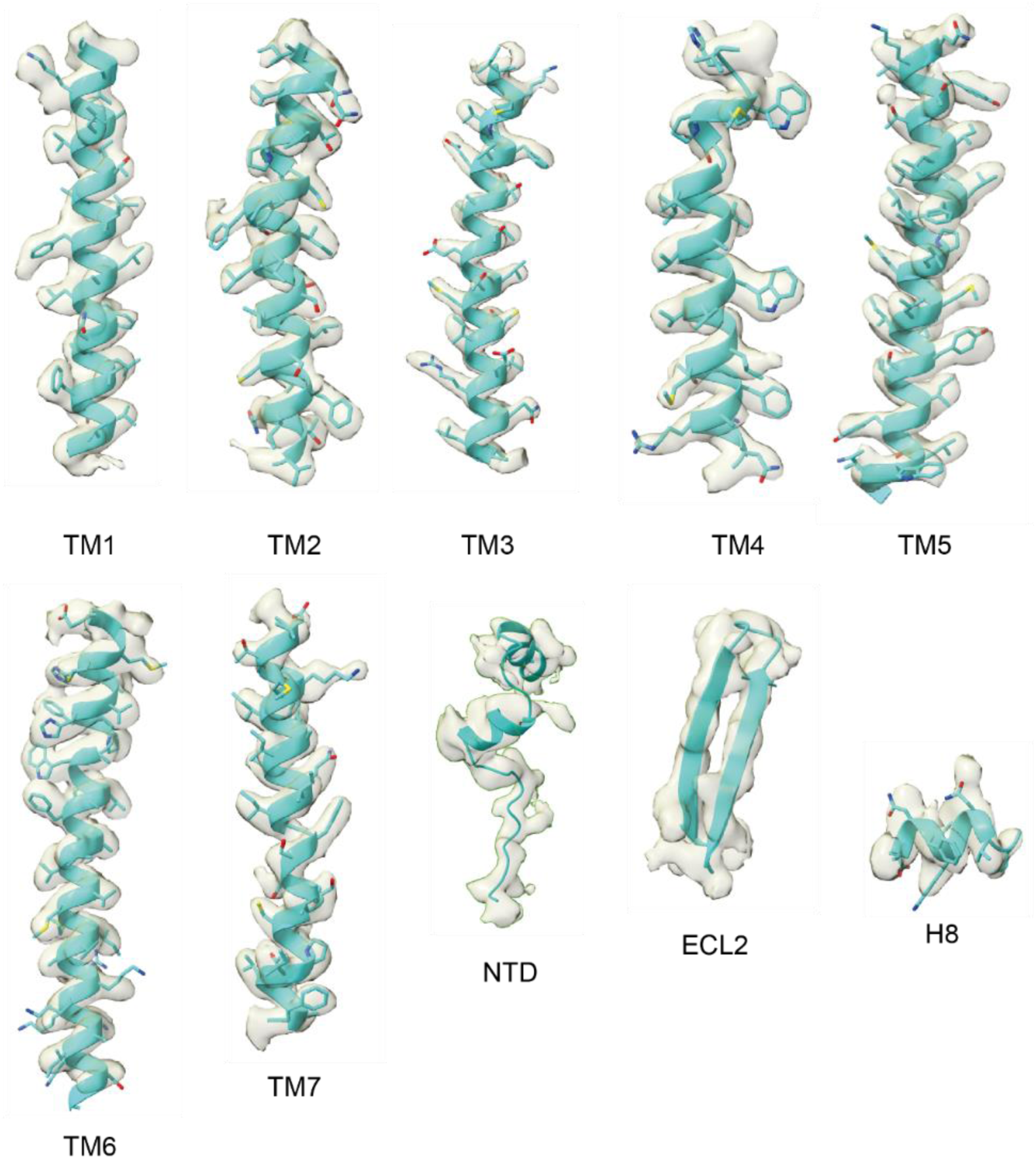
Representative densities of QRFPR in the 26RF-QRFPR-G_q_-scFV16 complex. Densities of seven transmembrane helices, helix 8, N-terminus, and ECL2 are shown.

**Supplementary Fig. S3.**
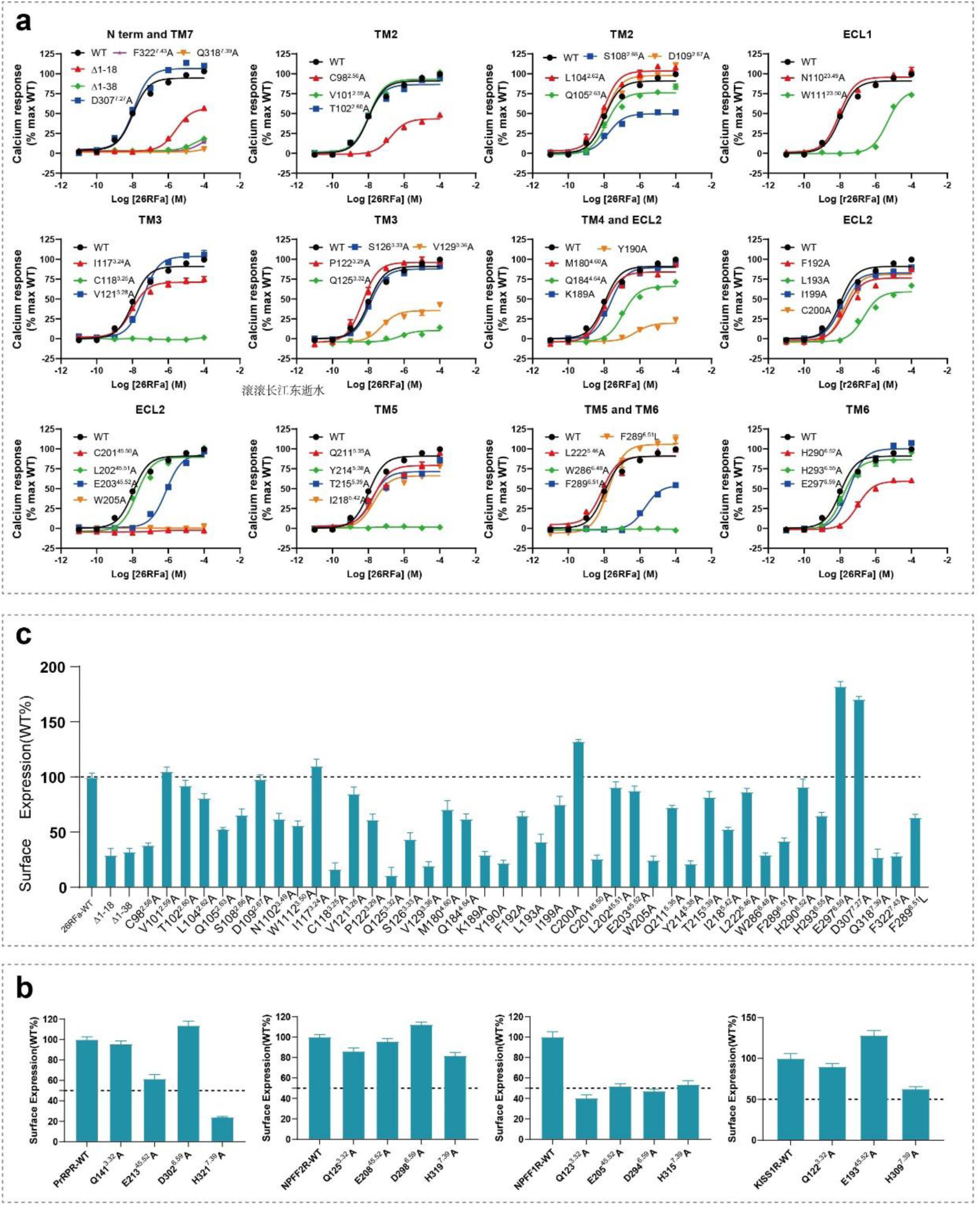
Effects of residue mutations in the QRFPR on 26RFa activity. **a**, Dose-response curves of 26RFa on mutants of QRFPR. **b,** Surface expression of QRFPR mutants. **c,** Surface expression of critical residues in the RF-amide binding residues.

**Supplementary Fig. S4.**
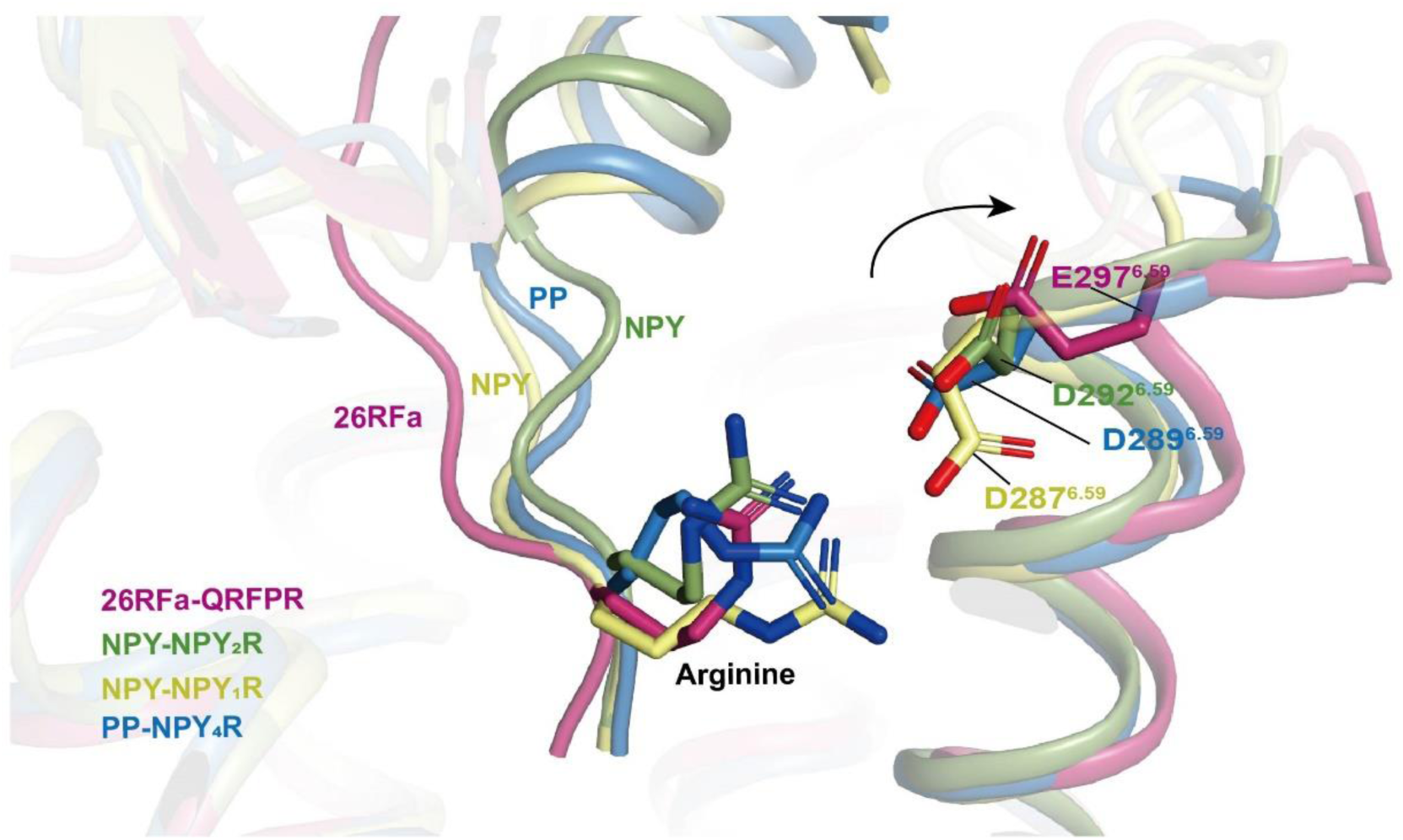
Comparison of interactions between the arginine in the RF/RY-amide motifs in 26RFa/NPY/PP and D/E^6,59^. The TM6 in QRFPR shows an outward movement relative to NPY bound NPY_1_R (PDB: 7X9A) and NPY_2_R (PDB: 7X9B), and PP bound NPY_4_R (PDB: 7X9C). Thus, in contrast to D^6,59^ in NPYRs, the residue E^6,59^ in QRFPR is positioned far from forming a salt bridge with the conserved arginine in the RF-amide motif.

**Supplementary Table S1.** Cryo-EM data collection, model refinement and validation statistics for the 26RFa-QRFPR-G_q_ protein complex.

| 26RFa-QRFPR-G<sub>q</sub> protein-scFV16 complex |  |
| --- | --- |
| Data collection and processing |  |
| Magnification | 165,000 |
| Voltage (kV) | 300 |
| Electron exposure (e<sup>-</sup>/Å<sup>2</sup>) | 50 |
| Defocus range (μm) | -0.8 ~ -1.8 |
| Pixel size (Å) | 0.73 |
| Symmetry imposed | C1 |
| Initial particle projections (no.) | 2,397,272 |
| Final particle projections (no.) | 202,884 |
| Map resolution (Å) | 2.73 |
| FSC threshold | 0.143 |
| Map resolution range (Å) | 2.0-4.0 |
| Refinement |  |
| Initial model used (PDB code) |  |
| Model resolution (Å, FSC=0.5) | 3.23 |
| Model composition |  |
| Non-hydrogen atoms | 9,543 |
| Protein residues | 1,208 |
| Ligand | 0 |
| Water | 0 |
| *B* factors (Å<sup>2</sup>, min/max/mean) |  |
| Protein | 30.00/126.95/74.69 |
| Ligand | -/-/- |
| Water | N/A |
| R.m.s. deviations |  |
| Bond lengths (Å) | 0.005 |
| Bond angles (°) | 0.979 |
| CC (mask) | 0.68 |
| Validation |  |
| MolProbity score | 1.24 |
| Clashscore | 4.62 |
| Rotamer outliers (%) | 0.10 |
| Ramachandran plot |  |
| Favored (%) | 98.99 |
| Allowed (%) | 1.01 |
| Disallowed (%) | 0.00 |

